# PLK1 O-GlcNAcylation is essential for dividing mammalian cells and inhibits uterine carcinoma

**DOI:** 10.1101/2022.08.21.504716

**Authors:** Sheng Yan, Bin Peng, Shifeng Kan, Guangcan Shao, Zhikai Xiahou, Xiangyan Tang, Yong-Xiang Chen, Meng-Qiu Dong, Xiao Liu, Xingzhi Xu, Jing Li

## Abstract

The O-linked N-acetylglucosamine (O-GlcNAc) transferase (OGT) mediates intracellular O-GlcNAcylation modification, whose function and substrates have entranced biologists and chemists alike. O-GlcNAcylation occurs on Ser/Thr residues and takes part in a vast array of physiological processes. OGT is essential for dividing mammalian cells, and it underscores many human diseases. Yet many of its fundamental substrates in the cell division process remains to be unveiled. Here we focus on its effect on Polo-like kinase 1 (PLK1), a mitotic master kinase that governs DNA replication, mitotic entry, chromosome segregation and mitotic exit. We found that PLK1 interacts with OGT and is O-GlcNAcylated. By utilizing stepped collisional energy/higher-energy collisional dissociation (sceHCD) mass spectrometry (MS) and mutagenesis studies, the critical O-GlcNAc site is located to be Thr291. Interestingly, T291N is a uterine carcinoma mutant in the TCGA database. Biochemical assays show that T291A and T291N both increase PLK1 stability. Using stable H2B-GFP cells, we show that PLK1-T291A and -T291N mutants display chromosome segregation defects, and result in misaligned and lagging chromosomes. In mouse xenograft models, we demonstrate that the O-GlcNAc-deficient PLK1-T291A and -T291N mutants would enhance uterine carcinoma in animals. Hence, we propose that OGT partially exerts its mitotic function through O-GlcNAcylation of PLK1, and sceHCD MS might be a new method to reveal many more O-GlcNAcylation substrates.

## Introduction

The O-linked N-acetylglucosamine (O-GlcNAc) glycosylation occurs on Ser/Thr residues and mediates cellular signal transduction by crosstalk with phosphorylation and ubiquitination (1,2). Currently about 5 000 proteins have been identified by proteomic or single-protein studies to be O-GlcNAcylated (3), and the list is ever growing. The O-GlcNAc transferase (OGT) has been found to regulate many biological processes, including transcription (4), the immune response (5), neurodegeneration (6), metabolism (7) and cancer (8). In the case of tumor biology, OGT and O-GlcNAc is upregulated in many cancer types, including bladder cancer, breast cancer, lung cancer, just to name a few (9).

Cell division is one of the fundamental biological processes, in which one mother cell will give rise to two daughter cells. And OGT depletion has long been found to impede the cell cycle progression (10). OGT overproduction would lead to chromosome bridges (11). Conversely, the cell division machinery also regulates OGT. For instance, OGT protein abundance decreases during mitotic onset (12), and the checkpoint kinase 1 (Chk1) phosphorylates OGT to stabilize OGT during cytokinesis (13). Therefore the reciprocal regulation between the cell cycle and OGT ensures the fidelity of mitosis (14) (15).

A proteomic study aiming to elucidate mitotic OGT substrates was carried out, and many O-GlcNAcylated proteins were found to function during spindle assembly and cytokinesis, the later stages of the cell cycle (11). And one of the important OGT interactors is polo-like kinase 1 (PLK1). PLK1 is a mitotic master kinase regulating DNA replication, mitotic onset, spindle assembly, centrosome disjunction, chromosome segregation, spindle checkpoint and cytokinesis (16,17). As a pivotal mitotic kinase, PLK1 is subject to a plethora of post-translational modifications (PTMs): its activation, symbolized by Thr210 phosphorylation, is mediated by the Aurora A kinase together with Bora (18); its inactivation, as indicated by Thr210 de-phosphorylation, is modulated by the myosin phosphatase targeting subunit 1 (MYPT1)-protein phosphatase 1cβ (PP1cβ) complex (19); it is phosphorylated at Tyr217, Tyr425 and Tyr445 by the nonreceptor tyrosine kinase c-ABL to promote protein stability and activity in cervical cancer (20); it is deubiquitinated by ubiquitin-specific peptidase 16 (USP16) to promote kinetochore localization for proper chromosome alignment in early mitosis (21); it is ubiquitinated at Lys492 by cullin 3 (CUL3)-KLHL22 and thus removed from kinetochores to achieve stable kinetochore-spindle attachment (22); it is mono-methylated at Lys209 and Lys413 by SETD6 to antagonize pThr210 and thus inactivated (23,24); it is di-methylated at Lys191 by SET7/9 in early mitosis for accurate kinetochore-microtubule dynamics (25); it is also subject to small ubiquitin-related modifier (SUMO) modification at Lys492, which results in its nuclear import and increased protein abundance (26); PLK1 is degraded by the anaphase-promoting complex/cyclosome (APC/C) during mitotic exit (27). Such a multi-layered regulation showcases the intricate biological network to regulate the correct localization, the rise and fall in protein abundance and timely activation and inactivation of PLK1, so that a successful mitosis will ensue. The role of PLK1 is not confined to mitosis, it is also implicated in cell motility, epithelial-to-mesenchymal transition and cancer. And many targeted therapies are being developed to treat PLK1-associated diseases (28).

In the OGT-centered mitotic screen, PLK1 was found to colocalize with a subset of OGT (11). OGT overexpression would reduce PLK1 transcripts and PLK1 protein abundance. But it does not alter PLK1-pThr210 levels or the localization pattern of PLK1 (11). Here we present evidence that PLK1 is O-GlcNAcylated. Through stepped collisional energy/higher-energy collisional dissociation (sceHCD) mass spectrometry (MS), we show that Thr291 is the main O-GlcNAcylation residue. Mutations of T291A or T291N will decrease PLK1 ubiquitination, resulting in increased PLK1 protein levels and mis-aligned or lagging chromosomes. Through TCGA and xenograft analysis, we further correlate our biochemical and cytological studies with pathological consequences: T291A and T291N mutations and PLK1 upregulation are conducive to uterine carcinoma. Our work suggests that the methodology of sceHCD MS could be utilized to explore many more potential OGT substrates, and PLK1 O-GlcNAcylation is essential for dividing mammalian cells.

## Results

### PLK1 is O-GlcNAcylated at Thr291

As Plk1 is such an important mitotic kinase, and PLK1 has been reported to colocalize with OGT (11), we wondered whether OGT could directly O-GlcNAcylate PLK1. To this end, we first examined the potential biochemical interaction between OGT and PLK1. As shown in Fig. 1A, cell extracts were immunoprecipitated (IPed) with anti-OGT antibodies and the immunoprecipitates were immunoblotted (IBed) with anti-PLK1 antibodies. The two proteins co-IP. We also examined the interaction between overproduced proteins (Fig. 1B). Cells were transfected with Flag-Plk1 and Myc-OGT plasmids, then the cellular lysates were IPed with anti-Myc antibodies and IBed with anti-Flag antibodies. Again, the overproduced proteins associate. Recombinant GST-OGT proteins were also utilized. Cells were transfected with Flag-PLK1 and the cellular extracts were incubated with GST-OGT proteins (Fig. 1C). And GST-OGT proteins could pulldown Flag-PLK1 (Fig. 1C). We then examined PLK1 O-GlcNAcylation. As shown in Fig. 1D, wild-type (WT) Flag-PLK1 shows RL2 (an O-GlcNAc antibody) staining in the IB. Then we enriched for O-GlcNAcylation with the treatment of OGA inhibitor Thiamet-G (TMG) together with glucose as described previously(29), and the RL2 band increased intensity significantly. These results suggest that PLK1 is O-GlcNAcylated.

**Fig. 1.**
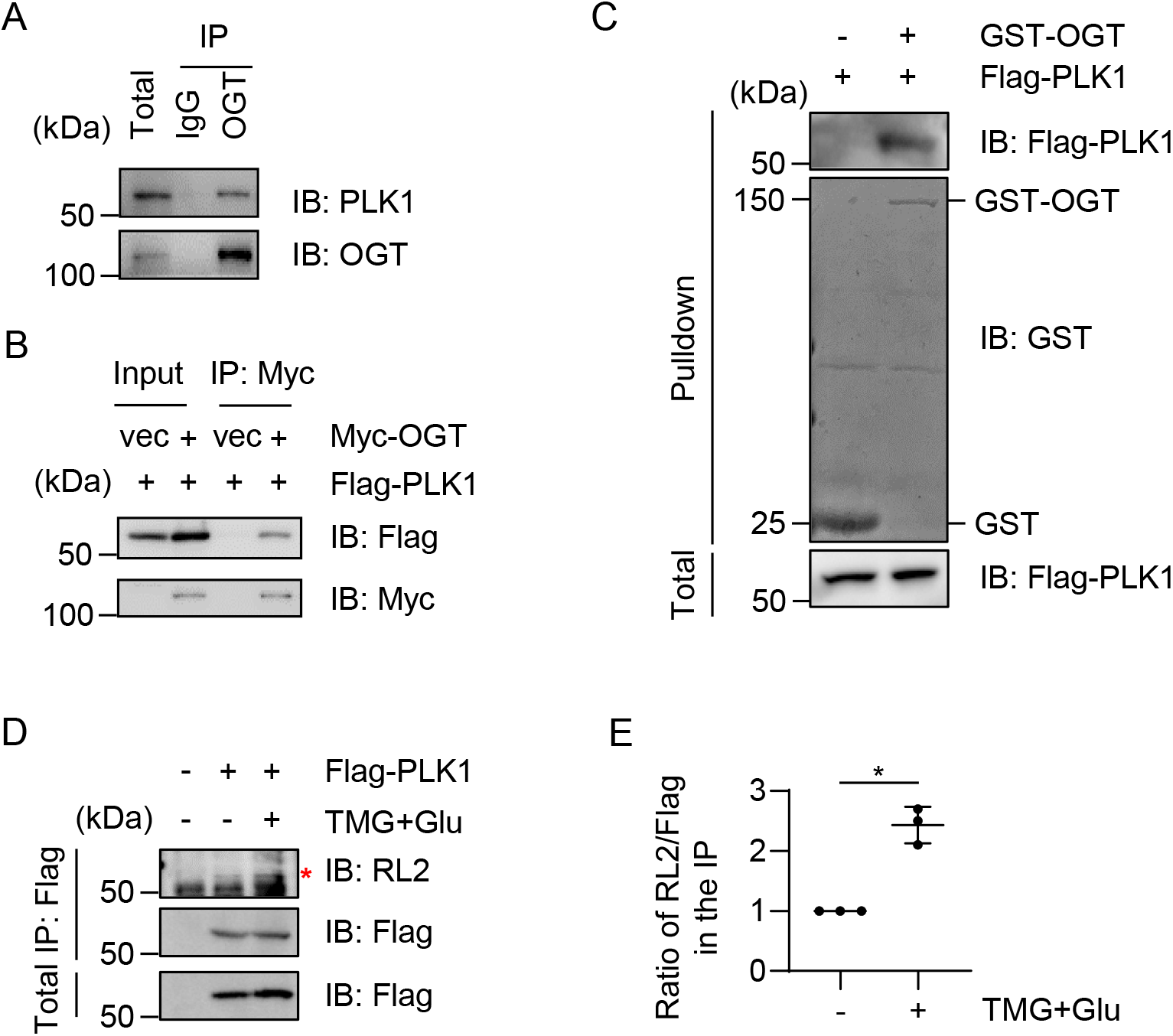
PLK1 is O-GlcNAcylated. (A) Endogenous PLK1 interacts with OGT. (B) Cells were transfected with Flag-PLK1 and Myc-OGT plasmids, and the cell lysates were immunoprecipitated and immunoblotted with the antibodies indicated. (C) Cells were transfected with Flag-PLK1. Recombinant GST-OGT proteins were incubated with the cellular lysates, and GST-pulldown experiments were carried out. (D) Cells were treated with OGA inhibitor Thiamet-G (TMG) plus glucose as previously described (29). And the cellular lysates were blotted with anti-O-GlcNAc antibodies RL2. Asterisk indicates the O-GlcNAcylated band. (E) Quantitation of (D). * indicates significant differences as determined by one-way ANOVA (p<0.05).

Then the Flag-PLK1-transfected cellular lysates were IPed, and the immunoprecipitates were subject to Mass spectrometry (MS) analysis (Fig. 2A). Electron-transfer dissociation (ETD) or higher-energy collisional dissociation (HCD) MS did not identify any sites (data not shown). Then we resort to sceHCD MS, as it is a method widely utilized for both N-glycopeptide analysis (30,31), and O-glycosylation characterization (32). sceHCD MS identified that the modification site could be Thr288 and Thr291 (Fig. 2A), and the number of spectra corresponding to Thr291 is three times more than Thr288. Therefore, we focused our study on Thr291. Incidentally, The Cancer Genome Atlas (TCGA) database reveals that T291N is associated with uterine serous carcinoma/uterine papillary serous carcinoma (cbioportal.org). So, we also included T291N in our study. Both T291A and T291N mutants greatly diminished RL2 signals (Fig. 2B). Sequence alignment shows that PLK1 Thr291 is conserved in various organisms, but not in fly or budding yeast (Fig. 2C). Taken together, our results suggest that PLK1 is O-GlcNAcylated and the major modification site could be Thr291.

**Fig. 2.**
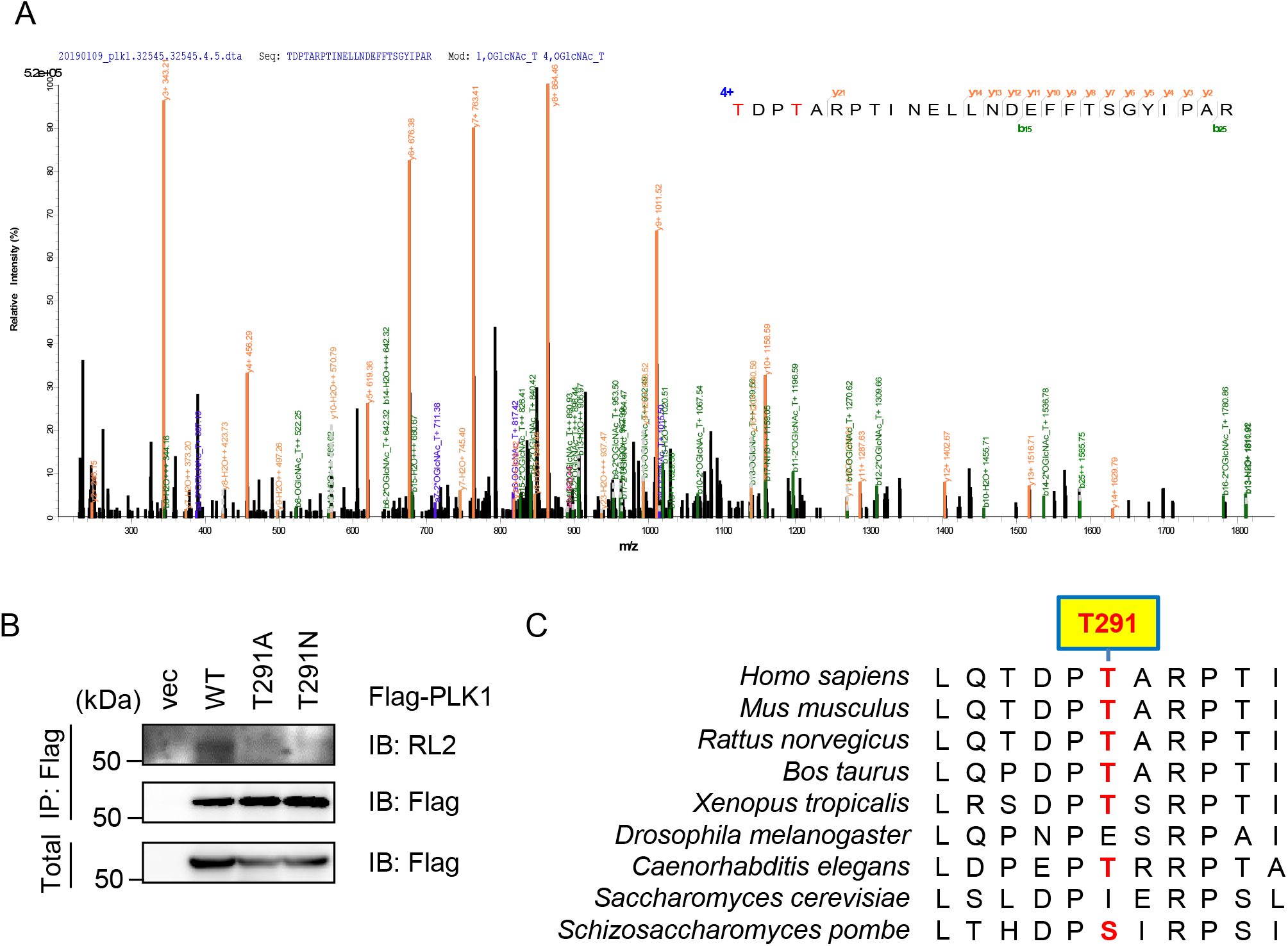
PLK1 is O-GlcNAcylated at Thr291. (A) Cells were transfected with Flag-PLK1, and enriched for O-GlcNAcylation. Anti-Flag immunoprecipitates were subject to Stepped collisional energy/higher-energy collisional dissociation (sceHCD) Mass Spectrometry analysis. And the result reveals that Thr291 could be an O-GlcNAcylation site. (B) Cells were transfected with PLK1-WT or -T291A, -T291N plasmids and the lysates were subject to immunoprecipitation and immunoblotting assays. (C) Thr291 of PLK1 is conserved in multiple species.

### PLK1 O-GlcNAcylation promotes ubiquitination

As the previous study suggests that OGT overproduction reduces PLK1 protein abundance (11), we therefore examined for PLK1 ubiquitination levels in the O-GlcNAcylation mutant. As shown in Fig. 3A, cells were transfected with Flag-PLK1, Myc-OGT and HA-Ub plasmids. And OGT overexpression markedly increased PLK1 ubiquitination levels. We also measured PLK1 ubiquitination by using the chemical OGT inhibitor Acetyl-5S-GlcNAc (5S) (Fig. 3B), and PLK1 ubiquitination levels decreased upon OGT inhibition. Then we used the O-GlcNAc-deficient mutant of T291A and T291N (Fig. 3C), and the mutants downregulated ubiquitination levels compared with the WT, suggesting that O-GlcNAcylation will increase PLK1 ubiquitination.

**Fig. 3.**
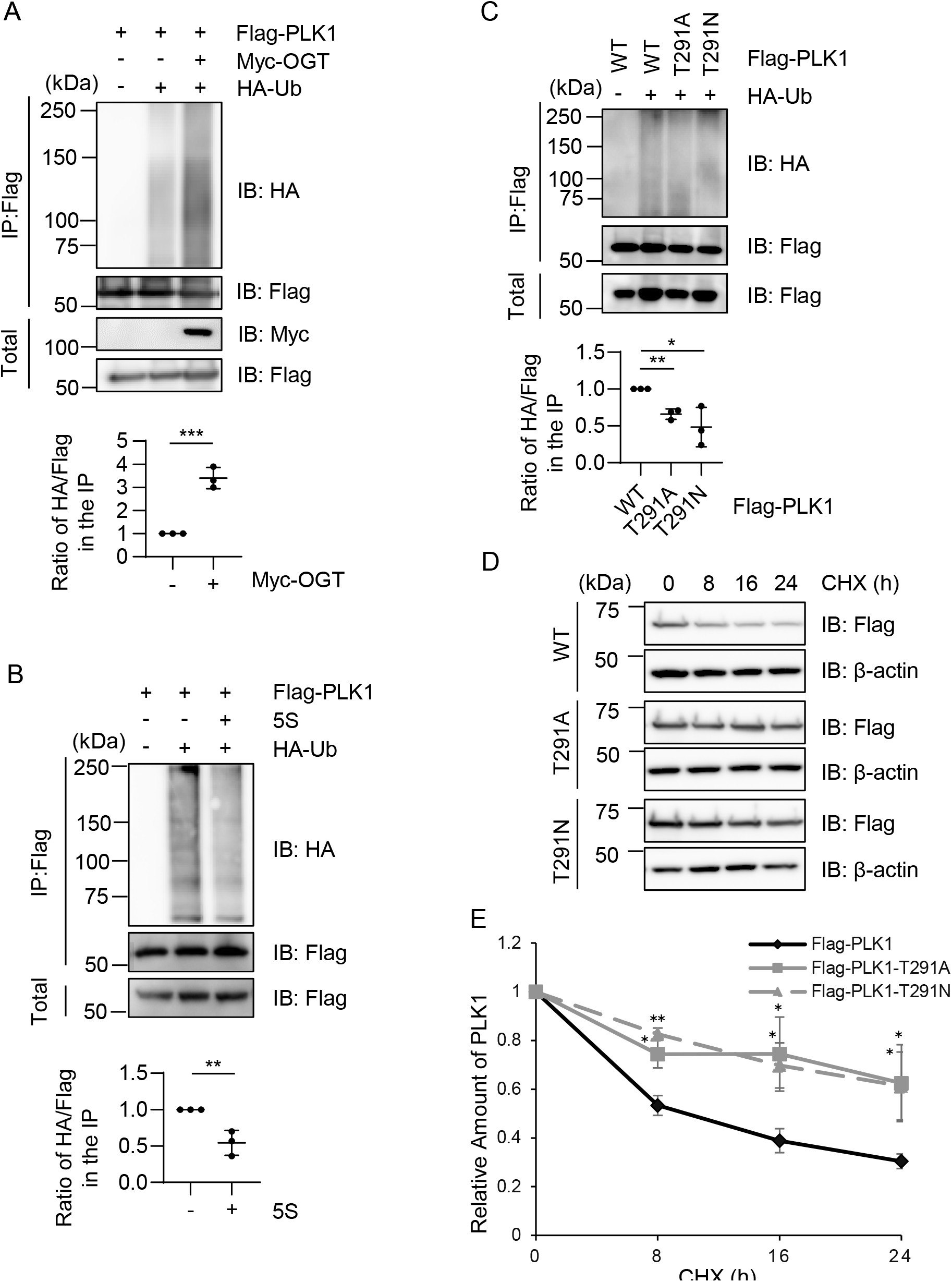
PLK1 O-GlcNAcylation promotes ubiquitination. (A) Cells were transfected with Flag-PLK1, Myc-OGT and HA-Ub. The lysates were immunoprecipitated with anti-FLAG antibodies and immunoblotted with the antibodies indicated. (B) Cells were transfected with Flag-PLK1 and HA-Ub, then treated with the OGT inhibitor Acetyl-5S-GlcNAc (5S) or not. (C) Cells were transfected with Flag-PLK1-WT, -T291A, -T291N and HA-Ub. Quantitation was carried out with one-way ANOVA, * indicates p<0.05; ** indicates p<0.01, *** indicates p<0.001. (D-E) Cycloheximide (CHX) pulse-chase assays. Cells were transfected with PLK1-WT, -T291A or -T291N, then treated with CHX for different durations. The quantitation is in (E). A two-way ANOVA test was used for statistical analysis. * indicates p<0.05; ** indicates p<0.01.

Then we employed the cycloheximide (CHX) pulse-chase experiments (Fig. 3D). Cells were transfected with Flag-PLK1-WT, -T291A, or -T291N plasmids, and treated with CHX to inhibit new protein synthesis. The cellular lysates were collected at different time points to examine for protein stability. As Fig. 3 D-E showed, the O-GlcNAc-deficient T291A and T291N mutants increased protein half-life compared to WT, consistent with the ubiquitination assays. In sum, consistent with previous investigations (11), our biochemical results show that O-GlcNAcylation de-stabilizes PLK1, probably through enhanced ubiquitination.

### PLK1 O-GlcNAcylation promotes mitotic progression

As PLK1 is an instrumental mitotic kinase (33), we assayed for mitotic defects of these O-GlcNAc mutants. We first constructed GFP-H2B HeLa cells stably expressing FLAG-vec, PLK1-WT, -T291A or -T291N (Fig. 4B), in which endogenous PLK1 is depleted by shRNA targeting PLK1 (Fig. 4C). When the cell cycle profile was assessed with flow cytometry, the T291A and T291N mutants manifested discernable increase in G2/M cells compared with the WT (Fig. 4 A-B).

**Fig. 4.**
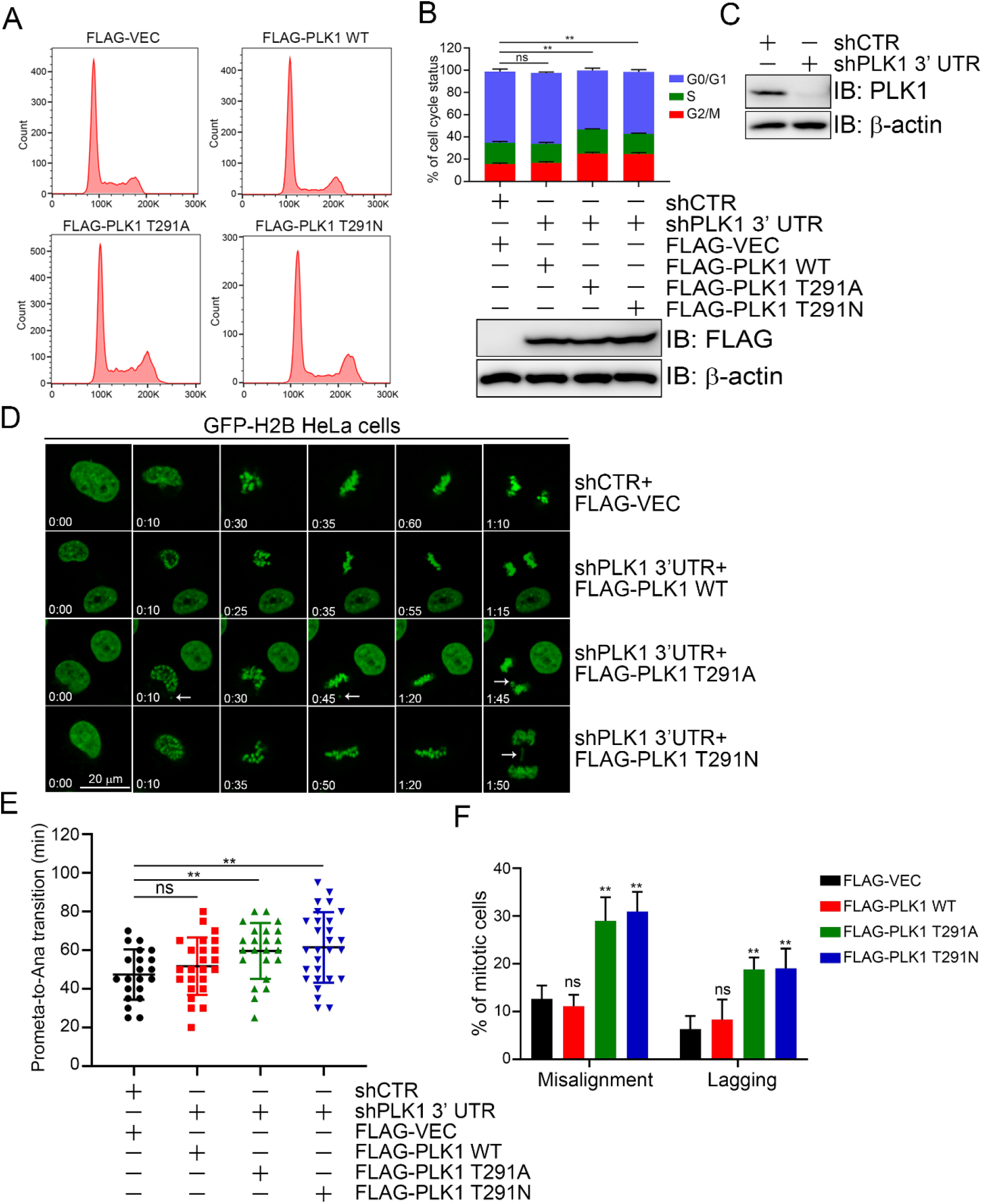
Both T291A and T291N of PLK1 delay mitotic progression, and induce chromosomal segregation defects. (A-C) T291A and T291N of PLK1 delay mitotic progression. GFP-H2B HeLa cells stably expressing FLAG-vec, PLK1-WT, -T291A or - T291N were constructed, where endogenous PLK1 was downregulated by shRNA targeting the 3’ UTR of *PLK1*. The expression of endogenous and exogenous PLK1 was analyzed by Western blots (B and C). Then the cells were stained with propidium iodide (PI) and analyzed by flow cytometry (A), and quantified in (B) (upper panel). (D-F) T291A and T291N of PLK1 induce chromosomal segregation defects. The cells in (A) were subject to time-lapse imaging. Representative time-lapse images are shown with the acquisition time relative to the onset of mitosis (D). The arrows indicate misaligned chromosomes or lagging chromosomes. The duration of prometaphase to anaphase onset in each group were quantified in (E). The percentage of misaligned or lagging chromosomes were quantified in (F). Scale bar, 20 μm. The data are presented as the mean ± SEM. Quantitation was carried out with the t test, **p < 0.01. ns, non-specific.

Then we analyzed chromosome segregation via time-lapse microscopy. The timing of cell cycle progression and phenotype of mitotic defects were monitored (Fig. 4D). Notably, the T291A and T291N mutants showed prolonged prometaphase to anaphase transition (Fig. 4E) and mis-aligned or lagging chromosomes (Fig. 4F). These defects are reminiscent of the SUMO mutant of PLK1 (K492R) (26). We examined for the possible crosstalk between PLK1 O-GlcNAcylation and SUMOylation, but no correlation was found (data not shown). We also measured for pT210, and the T291A and T291N mutants do not affect pT210 either (data not shown). These results indicate that PLK1 O-GlcNAcylation, and possibly its resultant PLK1 degradation, is essential for a faithful mitotic cell cycle.

### PLK1 O-GlcNAcylation inhibits uterine carcinoma

As T291N is a mutant involved in uterine carcinoma in the TCGA database, we investigated the correlation between PLK1 and uterine cancer. First, by bioinformatics analysis, PLK1 mRNA levels in UCEC (Uterine Corpus Endometrial Carcinoma) samples from TCGA database were examined and PLK1 overproduction is observed in cancer samples (Fig. 5A). When survival curves were analyzed, Kaplan-Maier survival curves of UCEC patients with high PLK1 expression levels show a poorer prognosis (Fig. 5B) (https://cistrome.shinyapps.io/timer/). Second, we constructed stable PLK1-WT, -T291A and -T291N HeLa cells (Fig. 5C), and the PLK1 expression levels were comparable. Third, the cell lines were injected into nude mice and tumor growth was monitored (Fig. 5D). The T291A or T291N mice exhibited increased tumor volume and growth compared to the WT (Fig. 5E-F), suggesting that chromosome mis-segregation and other mitotic defects might underlie uterine cancer growth.

**Fig. 5.**
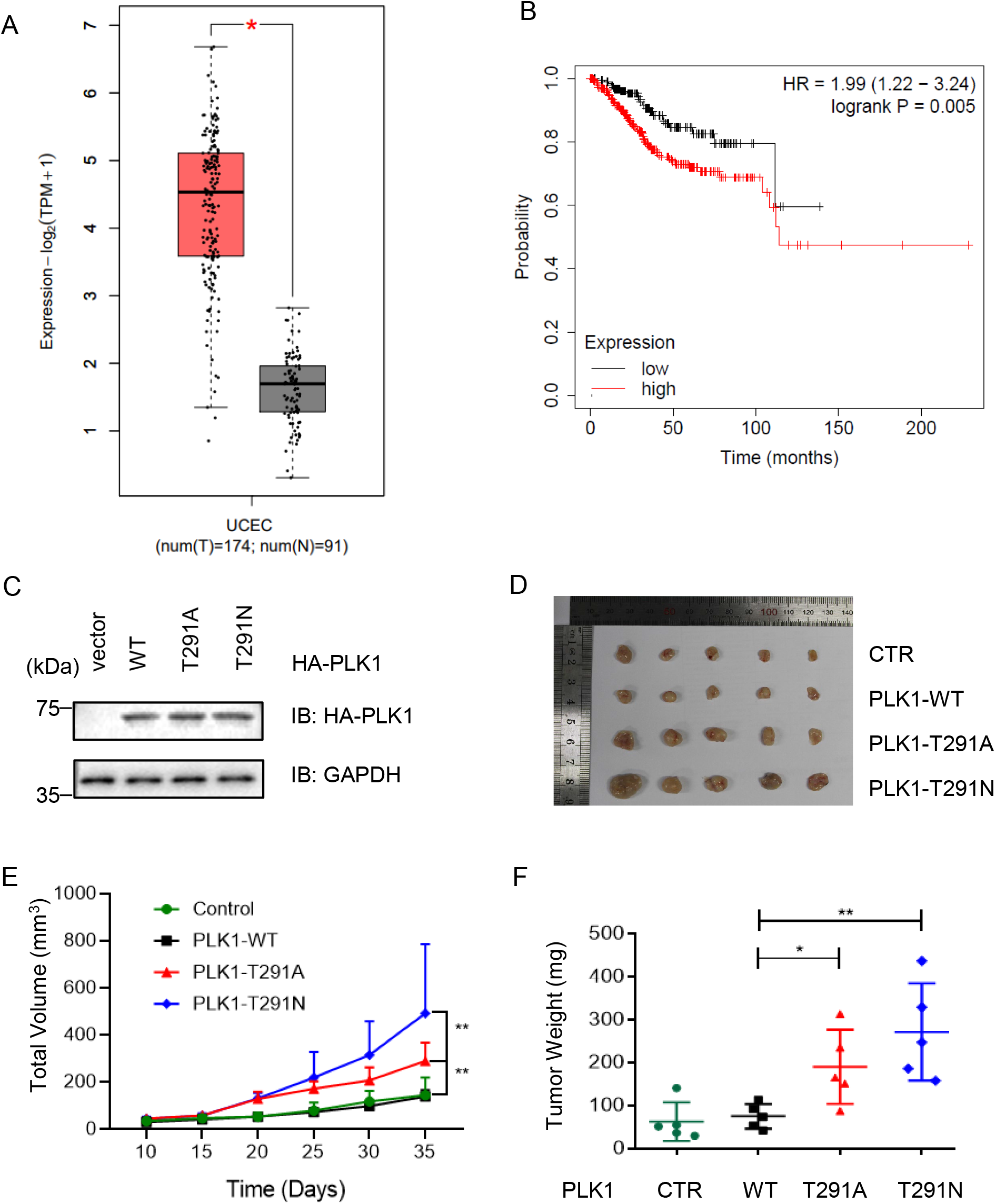
PLK1 O-GlcNAcylation mutants induce uterine carcinoma in xenograft models. (A) PLK1 mRNA levels in Uterine Corpus Endometrial Carcinoma (UCEC) samples from The Cancer Genome Atlas (TCGA) database. (B) Kaplan-Maier survival curves of UCEC patients with high or low PLK1 expression levels (https://cistrome.shinyapps.io/timer/). P-value was calculated with the chi-squire test. (C) Establishment of PLK1-WT and mutant stable cell lines in HeLa cells. (D-E) Xenografts in nude mice. PLK1-WT, -T291A, and - T291N cells were injected into nude mice. The tumors were photographed after 35 d. Tumor images were in (D), and tumor volumes were in (F). Quantitation was carried out with the t test, *p<0.05; **p<0.01.

## Discussion

In this paper we found that the mitotic master kinase PLK1 is O-GlcNAcylation, and further identified a major O-GlcNAcylation site to be Thr291 by sceHCD MS. Thr291 O-GlcNAcylation is instrumental for mitotic progression and the abrogation of which contributes to uterine carcinoma in xenograft models and perhaps in humans. Our findings are in line with previous observations that OGT partially colocalizes with PLK1, and negatively regulates PLK1 protein stability (11).

Previously sceHCD MS has been widely adopted in N-glycopeptide identification, and has been recently utilized in O-glycosylation analysis (32). O-GlcNAc site identification has been known to be challenging due to low stoichiometry, ion suppression of other peptides and its intrinsic feature of being easily hydrolyzed (1). Here by sceHCD MS, we found PLK1 O-GlcNAcylation sites. Through mutagenesis assays, Thr291 is found to be the major O-GlcNAc site. Our results suggest that sceHCD MS could be utilized for O-GlcNAc site mapping.

Many proteomic profiling work has shown a negative crosstalk between O-GlcNAcylation and ubiquitination (34), that is, O-GlcNAcylated proteins tend to be more stable. In the case of PLK1, however, our data show that O-GlcNAcylation promotes ubiquitination and subsequent degradation. We sought to identify the underlying mechanisms by examining the possible effect on the interaction between PLK1 and Cdh1 (also known as Fzr1), as PLK1 is degraded during mitotic exit by APC/C (27). But no alteration was observed in the O-GlcNAc-deficient T291A or T291N mutants. It is possible that O-GlcNAc might interfere with the association of PLK1 and other E3 ligases.

We also tried to examine the possible crosstalk between O-GlcNAc with other PTMs. Structurally, PLK1 comprises an N-terminal kinase domain (amino acid 53-303), a D-box (337-345) and a C-terminal Polo-box domain (345-603) (35). So Thr291 resides in the kinase domain. But T291A or T291N does not affect or the activation phosphorylation pThr210 (data not shown), consistent with previous results (11). SUMOylation at Lys492, another PTM that modulates PLK1 stability (26), was examined. And PLK1 O-GlcNAcylation does not crosstalk with SUMOylation either (data not shown), probably due to the long distance between the two PTM sites.

Investigators have long discovered that OGT depletion would hamper cell cycle progression, and many OGT substrates during the cell division have been identified (11) (36,37), suggesting that OGT plays various roles in mitosis. Our work here demonstrates that PLK1, the master mitotic kinase, is O-GlcNAcylation, and its abrogation would lead to mitotic defects. PLK1 is upregulated in many cancer types, and PLK1 inhibitors have been actively pursued as an anti-cancer therapy (38). As a “druggable target”, PLK1 inhibitors such as BI2536 and BI6727 (volasertib) were developed targeting its kinase domain, with volasertib reaching phase III trials (38). Our new finding of Thr291 O-GlcNAcylation revealed a new layer of regulation of PLK1, and may provide extra targets for designing next-generation inhibitors.

## Materials and Methods

### Cell culture, antibodies and plasmids

HeLa cells were purchased from ATCC. *OGT* plasmids and OGT antibodies were described before (13). PLK1 antibodies were Santa Cruz, sc-17783; RL2 antibodies were Abcam, AB2739. *PLK1* mutant plasmids were generated using specific primers (sequences available upon request) following the manufacturer’s instructions (QuickChange II, Stratagene). sh*PLK1* UTR: CCGGAGCTGCATCATCCTTGCAGGTCTCGAGACCTGCAAGGATGATGCAGCTTTTTT

### Immunoprecipitation and immunoblotting assays

immunoprecipitate and immunoblot experiments were performed as described before (39). The following primary antibodies were used for immunoblot: anti-HA (1:1000), and anti-FLAG M2 (Sigma) (1:1000), anti-Myc (1:1000), anti-PLK1 (1:1000). Peroxidase-conjugated secondary antibodies were from JacksonImmuno Research. The ECL detection system (Amersham) was used for immunoblot. LAS-4000 was employed to detect signals, and quantitated by the Multi Gauge software (Fujifilm). All western blots were repeated for at least three times. Chemical utilization: acetyl-5S-GlcNAc (5S-G) (OGT inhibitor) was used at 100 μM (prepared at 50 mM in DMSO) for 24 hrs.

### Stepped collisional energy/higher-energy collisional dissociation (sceHCD) Mass Spectrometry

For identification of O-GlcNAcylation by MS, PLK1 proteins isolated by gel electrophoresis were digested with trypsin (Promega) in 100 mM NH_4_HCO_3_ pH 8.0. The LC-MS/MS analysis was performed on an Easy-nLC 1000 II HPLC (Thermo Fisher Scientific) coupled to a Q-Exactive HF mass spectrometer (Thermo Fisher Scientific). Peptides were loaded on a pre-column (75 μm ID, 6 cm long, packed with ODS-AQ 10 μm, 120 Å beads from YMC Co., Ltd.) and further separated on an analytical column (75 μm ID, 12 cm long, packed with Luna C18 1.9 μm 100 Å resin from Welch Materials) using a linear gradient from 100% buffer A (0.1% formic acid in H_2_O) to 30% buffer B (0.1% formic acid in acetonitrile), 70% buffer A in 75 min at a flow rate of 200 nL/min. The top 20 most intense precursor ions from each full scan (resolution 120,000) were isolated for HCD MS2 {resolution 15,000; three-step normalized collision energy (25, 27, 30)} with a dynamic exclusion time of 60 s. Precursors with a charge state of 1+, 7+ or above, or unassigned, were excluded.

The software pFind 3 (40,41) was used to identify O-GlcNAcylated peptides by setting a variable modification of 203.0793 Da at S, T. The mass accuracy of precursor ions and that of fragment ions were both set at 20 ppm. The results were filtered by applying a 1% FDR cutoff at the peptide level and a minimum of one spectrum per peptide. The MS2 spectra were annotated using pLabel (42).

### Cell Cycle Profile Analysis

GFP-H2B stably expressing HeLa cells were harvested with trypsin, fixed in 75% ice-cold ethanol overnight, and stained with propidium iodide (PI) at room temperature for 30 min, followed by DNA content analysis using a CytoFLEX flow cytometer (Beckman Coulter). The cell cycle distribution was analyzed using FlowJo software.

### Time-Lapse Microscopy Assays

HeLa cells stably expressing GFP-H2B were infected with lentiviral plasmids carrying FLAG-VEC, PLK1 WT, T291A or T291N constructs. Endogenous PLK1 was knocked down by shRNA targeting the 3’ UTR on PLK1. For all time-lapse recordings, the culture dish was placed in a microincubator to maintain proper environmental conditions (37°C). All images were acquired using an Andor Dragonfly confocal microscope.

### Mouse Xenograft Analysis

For xenograft assays, 5 × 10^5^ WT or mutant cells were resuspended in Matrigel (Corning) and then injected into the flanks of nude mice (6-8 weeks old). The tumor volumes were measured every five days. At 35 days after the injection, tumors were dissected. The mice were obtained from the Beijing SPF Biotechnology Co., Ltd. [Certification NO. SCXK (Jing) 2019-0010]. All animal work procedures were approved by the Animal Care Committee of the Capital Normal University (Beijing, China).

## Data Availability Statement

The mass spectrometry proteomics data have been deposited to the ProteomeXchange Consortium (http://proteomecentral.proteomexchange.org) via the iProX partner repository (43) with the dataset identifier PXD036141.

## Abbreviations

(ETD): electron transfer dissociation
(sceHCD): Stepped collisional energy/higher-energy collisional dissociation
(MS): Mass spectrometry
(O-GlcNAc): O-linked β-N-acetylglucosamine
(OGT): O-GlcNAc transferase
(PTM): post-translational modification
(IP): Immunoprecipitation
(IB): Immunoblotting
(5S): Acetyl-5S-GlcNAc
(PLK1): polo-like kinase 1
(CHX): cycloheximide

## Acknowledgements

We thank Drs. Xing Chen (Peking Univ.) and Hai-Ning Du (Wuhan Univ.) for reagents and helpful discussions. J. L. is supported by the National Natural Science Foundation of China (NSFC) fund (31872720) and R & D Program of Beijing Municipal Education Commission (KZ202210028043).

